# Generalization in motor learning: learning bimanual coordination with one hand

**DOI:** 10.1101/2024.04.29.591705

**Authors:** Yiyu Wang, Madison M. Weinrich, Yuming Lei, David L. Wright, Milap Sandhu, John J. Buchanan, Deanna M. Kennedy

## Abstract

The ability to coordinate movements between the hands is crucial for many daily tasks. However, the precise mechanisms governing the storage and utilization of bimanual movement and the distinct contributions of each limb in this process are currently not fully understood. Two key questions persist: 1) How is the neural representation of bimanual coordination stored in the brain, and 2) How is the information governing bimanual coordination shared between hemispheres? In this investigation, we used a virtual partner (VP) to systematically address these issues by allowing the same coordination pattern (CP) to be acquired with unimanual and bimanual movements. More specifically, we used four experimental groups: unimanual (left, right) VP, bimanual, and control conditions. For each condition, retention and transfer tests were administered immediately and 6 hours after the initial practice. The control condition employed the same protocol as unimanual conditions without practice. As anticipated, performance after practice and during retention sessions indicated that all groups learned to perform the target CP. Furthermore, generalization from unimanual to bimanual occurred when the same type of visual feedback (VF) was provided. Interestingly, the absence of VF impaired motor generalization from unimanual to bimanual condition unless the participants initially practiced the task bimanually. Taken together, our results demonstrated that both limbs could access the memory representation of the CP. However, this globally shared representation appeared to be encoded in the visual-spatial domain. The conditions without VF underscored the importance of proprioception in forming a motor representation in intrinsic coordinates.

**NEW & NOTEWORTHY:** Conventional views on acquiring bimanual skills stress the need for simultaneous engagement of both hands. However, our study challenges this notion by demonstrating that the coordination pattern learned in unimanual conditions significantly boosts subsequent bimanual coordination—a novel approach to skill acquisition. Yet, this advantage diminishes without visual feedback, resulting in a breakdown of the intended bimanual coordination, highlighting the limitations of relying solely on unimanual practice.

## INTRODUCTION

The ability to coordinate movements between the hands is crucial for many daily tasks, from basic activities like tying shoelaces to more intricate actions such as playing a musical instrument or performing surgeries. Bimanual coordination refers to a particular spatial and temporal relationships between the left and right hands. Unraveling the mechanisms and neural processes that guide bimanual coordination is not just key to grasping human motor control but also carries important implications for rehabilitation, sports skill development, and designing robots.

Over the past few decades, numerous studies have explored the control mechanisms and neural processes related to coordinating movements with both hands. Various methods, such as behavioral analysis (1–5), electroencephalography (EEG) (6), Functional Magnetic Resonance Imaging (fMRI) (7, 8), mathematical modeling (9–11), and transcranial magnetic stimulation (TMS) (12, 13) have shed light on this area. Some studies suggest that in right-hand dominant individuals, the left hemisphere plays a crucial role in orchestrating bimanual coordination in a feedforward manner (6, 8, 14). For instance, EEG data reveals an asymmetry in cortico-cortical coherence direction during bimanual coordination tasks, with information flowing from the left hemisphere to the right hemisphere (6). This aligns with findings from behavioral evidence and mathematical modeling indicating stronger bimanual coupling from the right to the left hand during rhythmic coordination tasks (9, 14). In addition, TMS studies show increased motor-evoked potential (MEP) at the left primary motor cortex (M1) after practicing bimanual coordination tasks, with the MEP size linked to task complexity (12, 13). Similarly, neural imaging illustrates the significance of the left-lateralized parietal-to-premotor activation network in governing bimanual coordination (8). These findings favor the callosal access hypothesis that bimanual coordination is lateralized to the dominant hemisphere, resulting in an asymmetric contribution of the left and right hands in controlling movements (9, 14, 15).

Alternatively, the asymmetry in bimanual coordination may stem from independent control mechanisms associated with each hand, highlighted by the dynamic dominance hypothesis(16). Early research using repetitive tapping tasks showed distinct efficiency in movement control between two hands (17, 18). Sainburg and colleagues propose that the dominant hemisphere specializes in predictive control of discrete movement paths, while the non-dominant hemisphere excels at controlling movement impedance, based on experiments with both healthy individuals (16, 19–21) and patients with unilateral stroke (22–24). Evidence in support of the dynamic dominance hypothesis also comes from single limb rhythmic tasks that explored learning and transfer after physical and observational practice (25, 26). Considering the hypothesis of distinct control mechanisms, it is plausible that bimanual coordination emerges from intermingling and interference between the two hands, resulting in asymmetries in bimanual performance once the two hands are synchronized.

Despite advancements in understanding the control mechanisms and neural processes linked to bimanual coordination, questions about the precise mechanisms driving bimanual coordination behavior remain highly debated. Two key questions persist: 1) How is the neural representation of bimanual coordination stored in the brain? 2) How is the spatiotemporal information guiding bimanual coordination shared between the hemispheres? The interlimb transfer paradigm, commonly used in previous studies, serves as an approach to tackle these questions. Specifically, evaluating the ability of one hand to perform a movement after training the opposite hand in the same movement offers insights into the sharing of learned information between the left and right hemispheres (16, 20, 26–30).

Previous research using the interlimb transfer paradigm indicates that the neural representation of spatiotemporal patterns governing bimanual movements seems to be effector-independent, abstractly encoded in the central nervous system (8, 26, 31, 32). In most studies, coordination patterns were acquired in one limb (e.g., between elbow and wrist joints) followed by testing in the opposite limb or acquired on one side of the body (e.g., coordination between the left arm and the left leg) followed by testing on the opposite side (26, 31, 33). For example, a study using this paradigm revealed that the coordination pattern learned in unimanual conditions, practiced with a virtual partner (VP), facilitated subsequent bimanual coordination to achieve the same pattern (32). However, this research did not specifically address the control mechanisms associated with bimanual coordination by disassociating hemispheric contributions.

In the current study, we modified a prior method (32), in which participants were instructed to synchronize with a VP – an oscillating cursor by controlling another cursor at the target coordination pattern. Here, we integrated VP and hand movements into one cursor, superimposed on the Lissajous plot representing the target coordination pattern. The aim was to investigate the control mechanisms and potential neural processes involved in bimanual coordination. Based on previous investigations (26, 31–33), our hypothesis posited that both hands can access the neural representation of the 90° coordination pattern, facilitating subsequent bimanual coordination performance, irrespective of which hand is trained first. This perspective suggests that bimanual actions may be independent of hemispheric specificity in movement control, supporting the idea that bimanual coordination itself is lateralized.

Coordination patterns can be expressed relative to an intrinsic (the reference frame is muscle- or joint based) or extrinsic reference frame (the reference frame is external to the body) (34). For example, in-phase bimanual coordination of two index fingers in the horizontal plane can be described as mirror movement symmetry (extrinsic coordinate) or the simultaneous activation of homologous muscles (intrinsic coordinate). Given the challenge in perceiving (35) and performing (5) the 90° coordination pattern in both coordinate systems due to the competition with the intrinsic patterns (i.e., in-phase and anti-phase patterns), proficiency in performing the 90° coordination pattern may require extensive practice (5, 36), or it can be achieved in a few minutes using the Lissajous-cursor visual display (37–40). The cursor, representing the summed trajectory of two coordinated components, superimposed on the front of the Lissajous plot, is assumed to reduce incidental constraints (e.g., perceptual, cognitive, and environmental constraints) (41). This experimental arrangement facilitates a rapid improvement in performing the goal coordination pattern, providing an advantage for exploring behaviors and neural processes associated with complex bimanual coordination tasks (2, 4, 37, 39, 42).

In unimanual conditions, the cursor trajectory is determined by the cumulative integration of both the virtual partner (VP) and hand movements. Subsequently, either the left or right hand is tasked with synchronizing its motion with the constant-speed oscillation of the VP. Previous research posits that unimanual-VP coordination, facilitated through visual input, involves identical information variables essential for bimanual coordination (32, 43). Bimanual and unimanual-VP coordination are postulated to adhere to the same coordination dynamic process (32, 43, 44). Consequently, our hypothesis asserts that the 90° coordination pattern can be proficiently acquired through practice in both unimanual and bimanual conditions. Following initial skill acquisition, a bimanual coordination test, encompassing formats with and without a visual feedback display, was administered immediately and after a 6-hour delay. The results from these tests are expected to be consistent with earlier studies showing that performance in bimanual coordination remains stable even after a 6-hour lapse (13, 40), indicating that participants learned the task. Moreover, the absence of visual guidance underscores the robustness of the neural representation of the coordination pattern, facilitated by increased internal focus utilizing proprioceptive feedback (1, 40). Consequently, it is posited that reliance on visual cues in unimanual conditions may impede bimanual coordination performance when visual feedback is removed, while performance after the bimanual condition practice is expected to remain unaffected. Considering the distinct feedback information integral to learning the 90° coordination pattern in both unimanual and bimanual conditions, this study has the potential to elucidate the control mechanisms of bimanual coordination regarding extrinsic and intrinsic coordinate systems.

## MATERIALS AND METHODS

### Participants

A total of 51 undergraduate students (Mean age = ± 22.01, SD = ± 1.96; Male = 21, Female = 30) from the Department of Kinesiology and Sports Management at Texas A&M University volunteered to participate in the study. The Edinburgh Handedness Inventory was used to confirm that all participants were right-hand dominant (45). The Institutional Review Board at Texas A&M University approved the procedures, and participants provided written informed consent before participation in the study.

### Apparatus

Participants sat facing a computer screen with their left and right hands (palms facing down) placed comfortably on the table approximately shoulder width apart (Fig. 1A), and The experimental task required abduction-adduction motions of the index fingers on the horizontal plane. A customized finger apparatus for each hand allowed the index fingers to abduct and adduct comfortably without moving the other fingers on the horizontal plane (Fig. 1B). The index fingers were inserted into a slot connected to a potentiometer that recorded finger position. The potentiometer signal(s) were converted to voltage (AD convertor, NI USB-6210 Board, National Instruments Corp, Austin, TX, USA), sampled at 100 Hz, and stored on a PC for off-line analysis (MATLAB 2022a, MathWorks Inc.). Once the participants were comfortable with their posture, the index fingers were strapped to prevent sliding from the finger slot. The thumb and other unused fingers were fixed by finger barriers. A wood partition prevented participants from seeing their hands when performing the task. A movement template and a cursor (the Lissajous plot) representing the goal 90° pattern and finger motion (bimanual and unimanual) were displayed on a 27-inch monitor directly in front of the participants (Fig. 1C, D). The general task was to produce a 90° coordination pattern in a bimanual condition (BM), and in a unimanual left, or unimanual right condition (Fig. 1E). In the unimanual left condition, the 90° coordination pattern was performed by coordinating the left index finger with a virtual partner (VP) simulated by the computer software (VPL condition). In the unimanual right condition, the 90° coordination was performed by coordinating the right index finger with a virtual partner (VP) simulated by the computer software (VPR condition).

**Figure 1.**
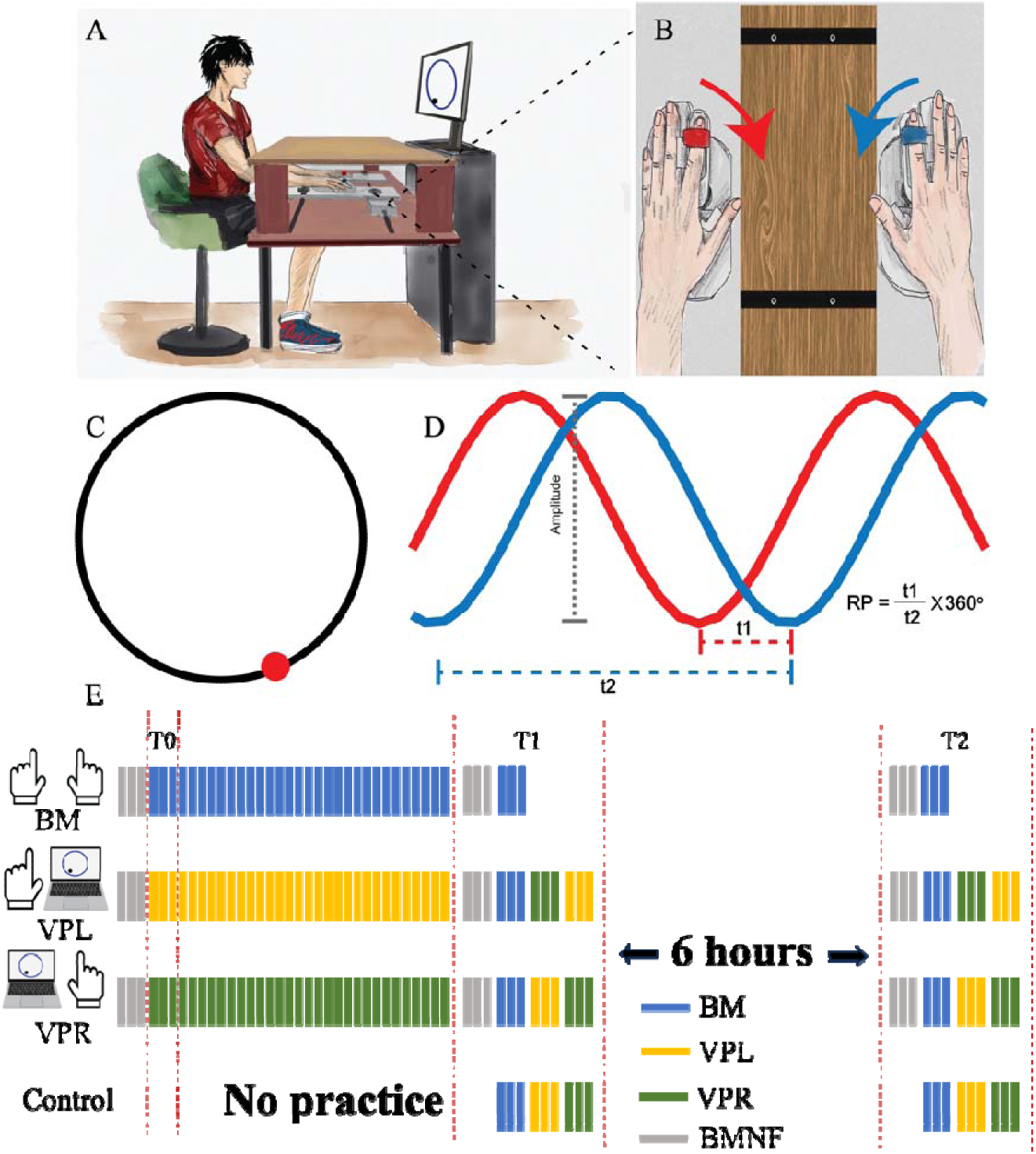
**(A)** Illustration depicting the experimental set up. **(B)** Hand and finger apparatus that measures finger abduction/adduction motion of the index fingers. **(C)** Illustration depicting the goal 90° relative phase Lissajous template and **(D)** corresponding example time-series to portray the calculation of relative phase (coordination). The t1 interval represents the time between consecutive left and right valleys, and t2 represents the time between valleys produced by the same limb. **(E)** Graphic representation of the experimental design and 4 experimental conditions: BM (blue), VPL condition (yellow), VPR condition (green), and control condition. All subjects received the same amount of practice trials except for the control group. The baseline (T0) was collected using the first three trials. The retention test was administered immediately (T1) and 6 hours after training sessions (T2). The gray bars represent the assessment of bimanual coordination with no visual feedback trials (BMNF).

### Procedure

Before entering the laboratory, participants were randomly assigned to one of four experimental conditions, including the BM (n = 12), VPL (n = 12), VPR (n = 14), and Control (n = 13) condition. Participants were instructed to rhythmically perform abduction-adduction movements on the horizontal plane with both index fingers in the BM condition or only move one finger if they were in the VPL or VPR condition.

A cursor on the screen was displayed by integrating the signals collected from the two potentiometers on the left and right sides for the BM condition (Fig. 1C). The cursor would move along the x-axis when the left index finger moved, and the cursor would move along the y-axis when the right index finger moved. In both VPL and VPR conditions, a cursor on the screen was displayed by integrating the signal collected from the left or right potentiometer with a computer-generated sine wave. The simulated sine wave moved the cursor at 0.6 Hz per cycle (A cycle is defined based on a cycle of sinewave from 0 to 2π). In the VPL and VPR conditions, the VP’s motion was plotted on the x-axis, while the finger motion was plotted on the y-axis. The participants were required to produce rhythmic movements by coordinating the left and right index fingers for the BM condition or coordinating one finger (left or right) with the VP for the VPL or VPR condition. The trained coordination pattern was a 90° relative phase between components, either the two fingers (BM) or a finger and the VP. The cursor and Lissajous template provided concurrent augmented feedback with respect to the 90° goal (Fig. 1C&D). Participants were instructed to move the cursor around the Lissajous plot to achieve the target 90° pattern in the BM, VPL, and VPR conditions. In the BM condition, participants were instructed to perform the 90° pattern at a self-selected frequency. Participants were reminded to increase the movement speed to make the coordination as rhythmic as possible without identifiable pauses during the trial (13, 37). In the VPL and VPR conditions, the simulated sinewave set the participant’s pace at 0.6 Hz as soon as the trial started. Participants were instructed to move either the left or the right index finger in a manner that would move the cursor around the Lissajous template. The same amount of practice was given in each experimental condition except for the Control condition, in which no practice trials were administered (Fig. 1E). The retention and transfer tests were administered for all conditions immediately (T1) after practice or after a 6-hour delay (T2). In BM, VPL, and VPR conditions, three bimanual coordination trials without the concurrent visual feedback (no cursor) (B_NF) were administered at the baseline (T0), and before two retest sessions (Fig. 1E).

Participants completed two lab visits for all four experimental conditions, BM, VPL, VPR, and control. The first visit was in the morning starting between 8:00 and 10:30 a.m. to avoid the influence of a daytime schedule on consolidation (46). Following the verbal instructions for the task, participants in the BM, BPL, and VPR conditions performed three B_NF trials as a measure of baseline performance. Each trial lasted 20 seconds with a 2-second break between trials. In the B_NF trials, participants coordinate their left and right hand at a self-selected pace while looking at the Lissajous template without the cursor providing concurrent movement status being displayed. Vision of the hands was blocked. The control group did not perform these trials. Following the B_NF trials at T0, participants produced thirty training trials (20 secs each trial, 2-sec break between) with the 90° coordination task in the BM, VPL, and VPR conditions. During training concurrent augmented feedback was provided with the cursor-Lissajous display. Participants were asked to move the cursor around the Lissajous template as accurately as possible. Following the practice trials, two retest sessions were administered, immediately after training (T1) and after a 6-hour delay (T2). In the retest sessions, participants in BM, VPL, and VPR conditions performed three trials of B_NF and three trials of BM to assess motor memory (Fig. 1E). Each trial was 20 seconds long with an inter-trial interval of 2 seconds. All required experimental procedures were completed after the test at T2. Participants in the BM task were finished after retest of the BM pattern at in session T2. Individuals in the VPL and VPR conditions were tested for each unimanual training condition after the bimanual test. Specifically, participants received three trials of VPL and three trials of VPR at T1 and T2 following the initial training. If participants practiced VPL first, testing VPR was denoted as a transfer R/VPL condition. If participants practiced VPR first, testing VPL was denoted as a transfer L/VPR condition. The duration of each trial was 20 seconds, and the inter-trial interval was 2 seconds. All required experimental procedures were completed after the retention test at T2.

Previous research demonstrated that cursor-Lissajous feedback significantly enhances performance in bimanual coordination tasks with a few minutes of practice (37, 38). It is plausible that brief exposure to bimanual practice guided by augmented visual feedback could enable the production of the 90° bimanual coordination pattern. To ensure that any benefits observed in bimanual performance were not solely due to a brief exposure to bimanual practice during retest sessions, a control condition was implemented. Participants in the control condition did not train on the 90° pattern and were first exposed in the retest sessions (Fig. 1E). B_NF trials were not included in the control condition. Bimanual coordination assessed in the control condition was labeled BC, the visual perceptual learning in the control condition as LC, and the visual perceptual retention in the control condition as RC.

### Measures and data reduction

All data computation and reduction were processed using MATLAB (R2022a). A low-pass Butterworth filter (10 Hz cutoff) was applied to the recorded potentiometer time series representing finger abduction-adduction motions. The filtered time-series were detrended to identify zero-crossing points to partition the time-series into half cycle segments, which were normalized to a range from −1 to 1.The absolute error (AE) of relative phase (ϕ) was calculated to determine the spatiotemporal accuracy of coordinated movements at the target goal of 90°. Relative phase was computed for each cycle of performance using the point-estimate approach (5). The relative phase was equal to the ratio of the time difference between the peak of signal one and the adjacent peak of signal two divided by the time difference between the same peak of signal one and the next peak of signal one (Figure 1D). The ratio was multiplied by 360°. The relative phase was computed between peaks based on the positive values and the negative values. The difference between each relative phase and the goal relative phase value (90°) was calculated to determine AE for each cycle of performance. In the end, all AEs were averaged to determine the spatiotemporal accuracy for a trial. In addition, the relative phase standard deviation (ϕSD) of the mean relative phase for each trial was computed to measure the stability of coordination performance.

### Statistics

We first performed a repeated measure ANOVA to assess motor performance (AE and ϕSD) improvement over practice with a within-group factor Trial (36 trials, including 30 practice trials and 6 retention trials) and a between-group factor Condition (BM, VPL, and VPR).

The motor performance (AE and ϕSD) of the same task from each retention and transfer test was calculated as the averaged performance from three trials. The RMANOVA was run to assess whether motor performance (AE and ϕSD) of the opposite hand benefited from the training of the right hand with a within-group factor Time (T1, T2) × a between-group factor Condition (L/VPR, LC), and whether the motor performance of the opposite hand benefited from the training of the left hand with a within-group factor Time (T1, T2) × a between-group factor Condition (R/VPL, RC). In addition, the BM performance with and without visual feedback was examined for each condition during the retention or transfer tests. The RMANOVA was run to assess and compare the BM performance with visual feedback after the practice for each condition with a within-group factor Time Time (T1, T2) × a between-group factor Condition (BM, BC, VPL, and VPR). Further, the RMANOVA was run to assess and compare the BM performance without visual feedback after the practice for each condition with a within-group factor Time Time (T1, T2) × a between-group factor Condition (BM, B_NF, VPL, and VPR). Hereby, BM, as a bimanual performance with visual feedback, was set as a controlling factor compared to BMNF performance for three other conditions. The result could also indicate the difference in BM performance between visual feedback and no visual feedback conditions.

For the repeated measure ANOVA test, the Greenhouse-Geisser adjusted degree of freedom and effect size was reported if the assumption of Sphericity was violated. Post hoc comparisons were performed using a Bonferroni test (α = 0.05) when appropriate. A power analysis (Gpower V3.1) indicated that 8 participants per group would be required for 80% statistical power given the reported effect size from the pilot result (partial η^2^ = 0.32) for the repeated measures ANOVA when only two groups were included in the analysis, while 5 participants per group would be required for achieving 80% statistical power when four groups were included in the analysis. The current experiment recruited 51 participants, which would provide sufficient power to support experimental findings.

## RESULTS

### Training

For all three groups, BM, VPL, and VPR, significant improvements emerged over the training interval in terms of performance error and coordination stability (Figure 2A-I). The analysis of the performance accuracy data (AE) revealed an improvement at the target pattern of 90° over the practice trials [, F (10.45, 365.79) = 6.37, *p* < 0.01, partial η^2^ = 0.15] (Figure 2J). The analysis of the AE data also revealed a significant main effect for Condition [F (2, 35) = 5.43, *p* = 0.01, partial η^2^ = 0.24], with *post hoc* analysis indicating that performance accuracy in the BM condition [mean ± SD: 33.276 ± 6.03] was significantly better than in the VPL condition [41.39 ± 6.03, *p* = 0.01], but not significantly different between the BM and VPR conditions, and the VPL and VPR conditions.

**Figure 2.**
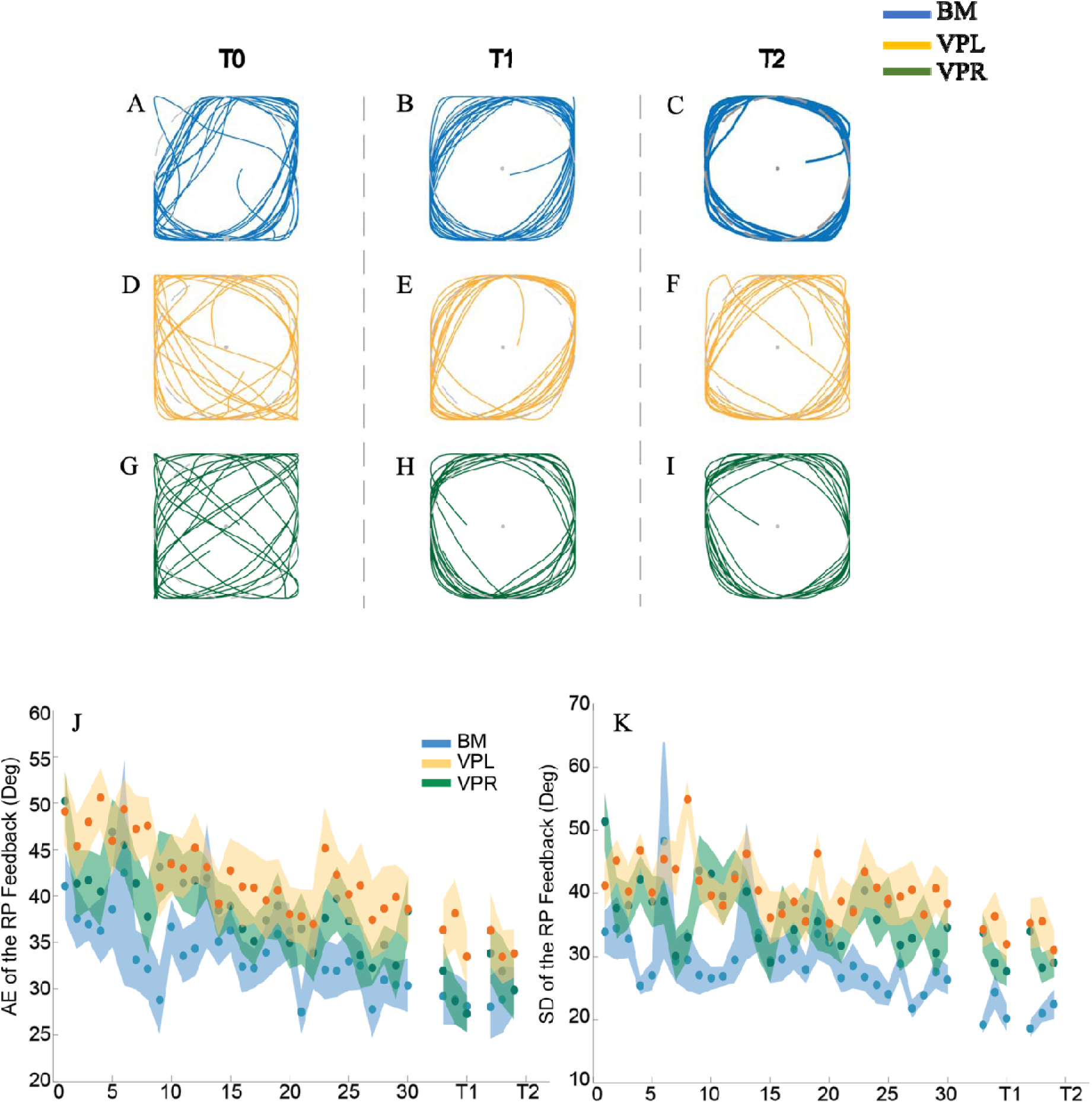
These graphs demonstrate performance improvement across training. (A-I) Sample performance for each condition at each testing blocks (T0, T1, T2) is presented. The BM condition (A-C), VPL condition (D-F), and VPR (G-I). (J) Graphic representation of absolute error (AE) of relative phase (coordination) in degrees across trials for the BM, VPL, and VPR groups. (K) The standard deviation (SD) of coordination in degrees (Deg).

The analysis of relative phase ϕSD data, the stability measure, indicated coordination stability significantly improved as a function of practice [F (8.31, 290.68) = 4.30, *p* < 0.01, partial η^2^ = 0.11] (Figure 2K). There was a significant Condition effect [F (2, 35) = 9.69, *p* < 0.01, partial η^2^ = 0.36], and post hoc analysis indicated the BM [28.20 ± 6.65] performance was less variable than the VPL [39.88 ± 6.65, *p* < 0.01] and VPR conditions [36.15 ± 6.65, *p* = 0.01]; while variability did not differ between the VPL and VPR conditions [p = 0.49]. Further, there was a significant interaction effect of Trial × Condition [F (70, 1225) = 1.43, p = 0.01, partial η^2^ = 0.08], suggesting variability in the BM condition was not always less than the VPL and VPR conditions. For example, there was no difference between the BM and VPL [*p* = 0.32], and the BM and VPR condition [*p* = 0.39] at Trial 22, while the BM condition was significantly more stable than the VPL and VPR conditions at Trial 25.

### Unimanual Transfer

Transfer for the unimanual conditions compared motor performance between L/VPR and LC, and R/VPL and RC to determine whether the untrained hand benefited from practice. The analysis detected a significant main effect of Condition indicating that overall accuracy from T1 and T2 was significantly greater for L/VPR than for LC [F (1, 25) = 11.49, *p* < 0.01, partial η^2^ = 0.32] (Figure 3A). However, there was no significant main effect for Time [F (1, 25) = 2.99, *p* = 0.10] and the Time × Condition interaction was not significant [F (1, 25) = 0.24, *p* = 0.63]. In addition, overall variability at T1 and T2 was greater for L/VPR than for LC [F (1, 25) = 21.62, *p* < 0.01, partial η^2^ = 0.46] (Figure 3B). While the analysis failed to detect the main effect of Time [F (1, 25) = 0.15, *p* = 0.70] and the Time × Condition interaction was not significant [F (1, 25) = 2.82, *p* = 0.11].

**Figure 3.**
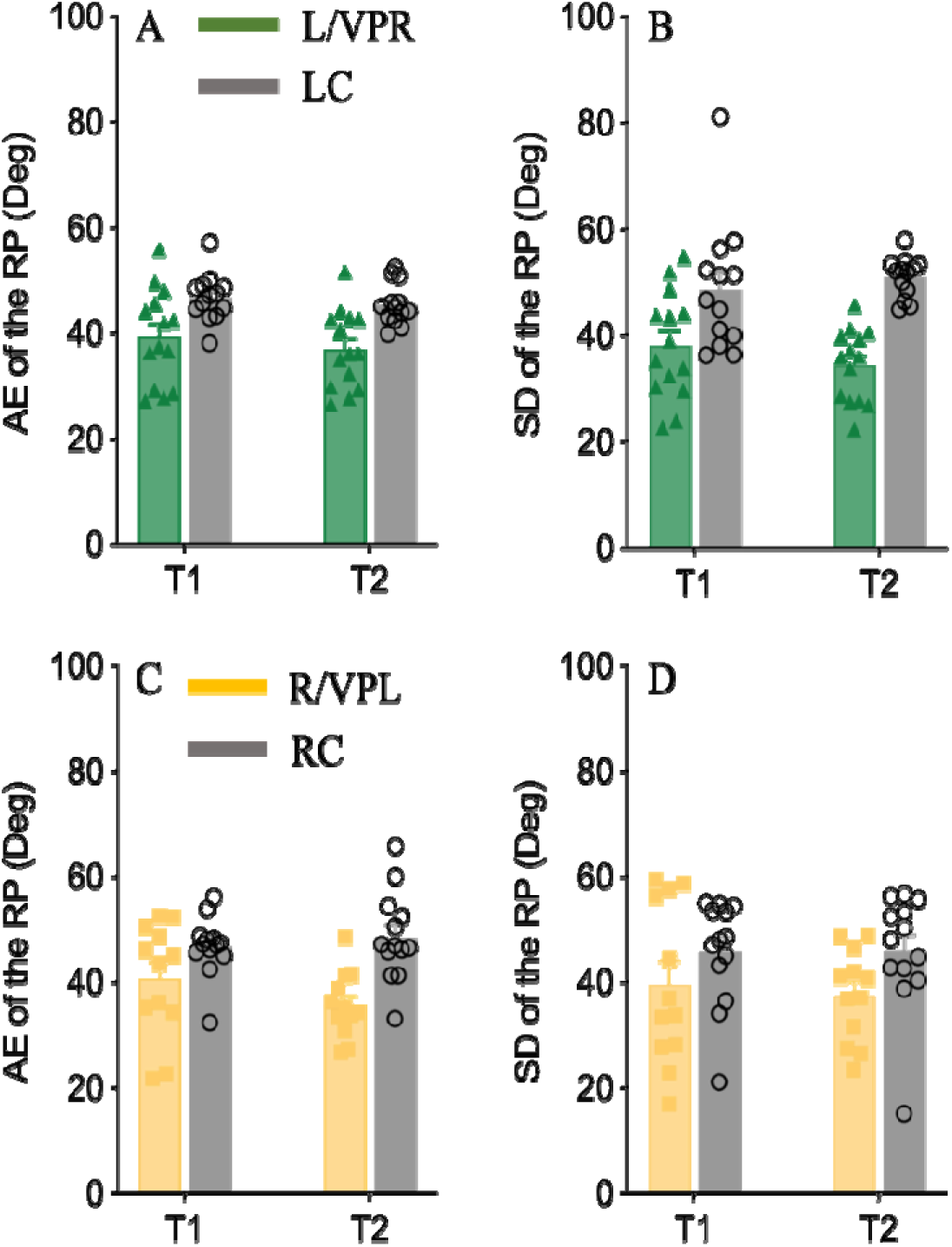
Generalization of learning from the trained limb to the untrained limb. AE (A) and SD measures of relative phase (B) are plotted for left limb transfer for the trained VPR group (L/VPR, green, closed triangles) and for controls with no training with a virtual partner using their left limb (LC, gray, open circles). (C and D) like the top row except the data represents performance of subjects using their Right limb after training VPL (R/VPL, yellow rectangles) compared to controls who had no training experience with the Right hand to coordinate with VP (RC, gray open circles). Error bars indicate standard error of the mean.

By comparing R/VPL and RC, the analysis detected a main effect of Condition indicating that overall accuracy for R/VPL was significantly greater than RC [F (1, 23) = 16.20, *p* < 0.01, partial η^2^ = 0.41] (Figure 3C). However, the analysis did not find a main effect for Time [F (1, 23) = 0.58, *p* = 0.45] or a significant Time × Condition interaction [F (1, 23) = 2.52, *p* = 0.13]. The analysis of the variability data indicated that the main effect of Condition for coordination stability approached the standard level of acceptable significance [F (1, 23) = 4.24, *p* = 0.05, partial η^2^ = 0.16], with R/VPL [38.60 ± 8.80] showing more variability than RC [45.85 ± 8.45] (Figure 3D). The analysis did not reveal a main effect for Time [F (1, 23) = 0.10, *p* = 0.75] and the Time × Condition interaction was not significant [F (1, 23) = 0.15, *p* = 0.71].

### Unimanual to Bimanual Transfer with visual feedback

To test for unimanual to bimanual transfer with visual feedback, the AE and ϕSD data from the visual feedback trials at T1 and T2 were compared across the four conditions, BM, BC, VPL, and VPR. The statistical analysis revealed a significant main effect of Condition [F (3, 47) = 9.18, *p* < 0.01, partial η^2^ = 0.37] (Figure 4A). The *post hoc* statistics further indicated that bimanual performance from the BM, VPL, and VPR conditions were significantly more accurate than the BC [*p* < 0.01] condition. In addition, the analysis indicated participants achieved similar levels of bimanual accuracy immediately after the BM, VPL, and VPR practice and in the 6-hour delayed test, with no difference between BM and VPL and BM and VPR [*p* = 1.00]. The analysis also detected a significant main effect of Time [F (1, 47) = 5.44, *p* = 0.02, partial η^2^ = 0.10], with the averaged accuracy from all conditions being significantly better at T2 [34.58 ± 12.17] than T1 [37.91 ± 14.17] (Figure 4A). The Time × Condition interaction was not significant [F (3, 47) = 1.00, *p* = 0.40].

**Figure 4.**
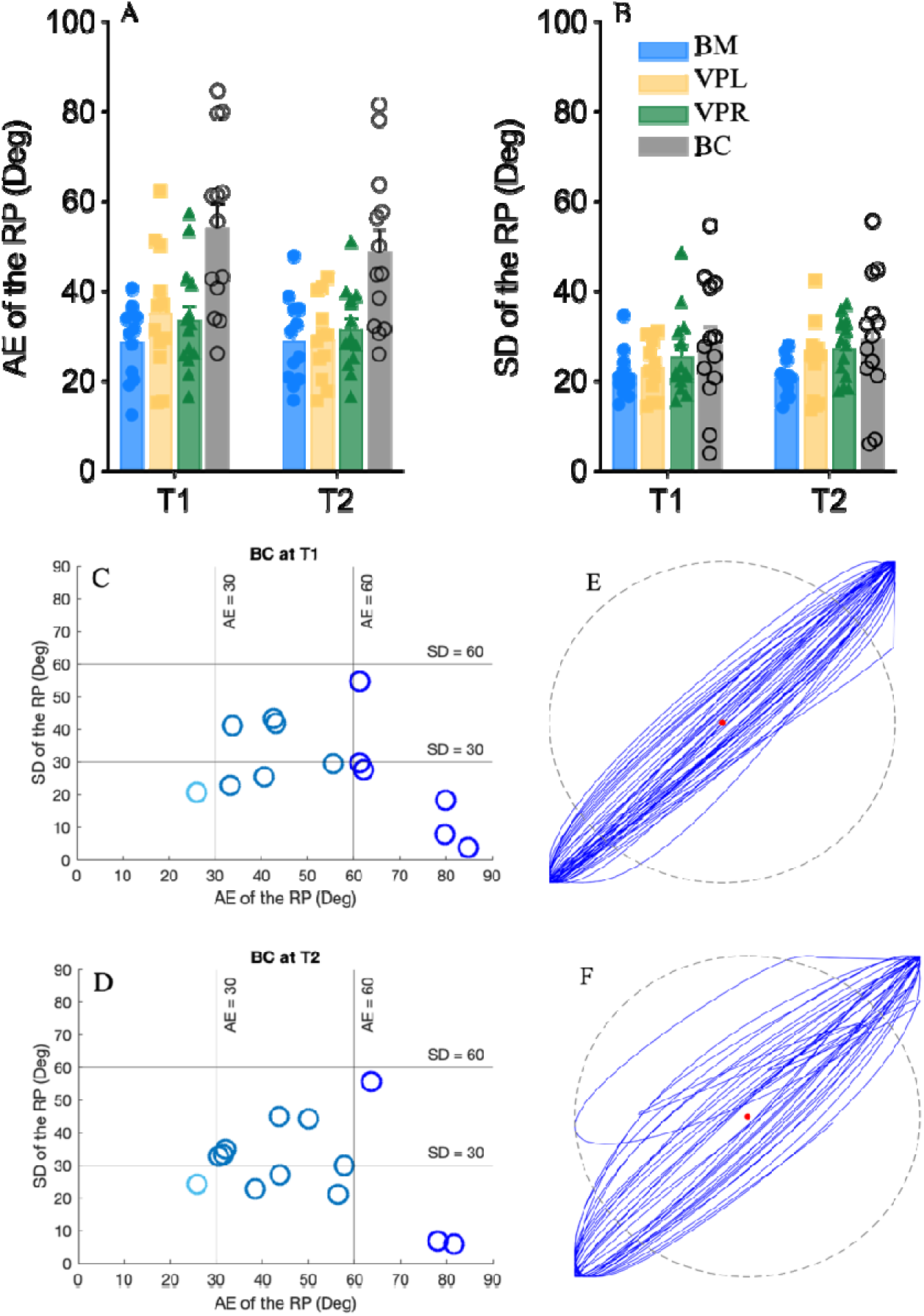
Depicts the mean AE (A) and SD (B) in degrees for each experimental condition at T1 and T2: bimanual training (BM, blue circles), VPL condition (yellow squares), VPR (green triangles), and the bimanual control (BC, open circle). Error bars represent the standard error of the mean. (C &D). (E &F) Examples illustrating attraction to in-phase coordination (T1 and T2), a predominant bimanual coordination pattern in the BC condition at T1 and T2. The XY coordinate is segmented into nine areas with the bottom-left corner (low AE and SD) indicating best performance. Despite the lower SD values observed in the BC condition, the larger AE indicates that performance was not stabilized at 90°.

The analysis of ϕSD failed to detect a main effect of Condition [F (3, 47) = 2.16, *p* = 0.11], or Time [F (1, 47) = 0.50, *p* = 0.48], and the Time × Condition interaction was not significant [F (3, 47) = 0.16, *p* = 0.93] (Figure 4B). Combining AE and ϕSD, the results indicate that individuals from the bimanual control condition (BC) with AE closer to 90° tended to perform with less variability. Given that the 0° and 180° coordination patterns are considered as intrinsic modes in the central nervous system, the plots in Figure 4C-F show that individuals in the BC condition performed with smaller variability due to an attraction to 0 ° and 180 ° instead of towards the required 90 pattern.

### Unimanual to Bimanual Transfer without visual feedback

To test for unimanual to bimanual transfer without visual feedback, the AE and ϕSD data from the visual feedback trials at T1 and T2 were compared across the four conditions, BM, B_NF, VPL, and VPR. (Figure 5A-L). The statistical analysis of AE revealed a main effect of Condition [F (3, 46) = 10.42, *p* < 0.01, partial η^2^ = 0.41] a main effect of Time [F (1.75, 80.68) = 12.43, *p* < 0.01, partial η^2^ = 0.21], and a significant Time × Condition interaction [F (6, 92) = 2.67, *p* = 0.02, partial η^2^ = 0.15] (Figure 5M). The post hoc analysis of the Time × Condition interaction revealed improved accuracy for B_NF at T1 and T2 [*p* < 0.05] compared to T0 as a function of practice, but the difference between T1 and T2 was not significant [*p* = 0.85]. At T0, accuracy in the B_NF condition was significantly greater than VPR [p < 0.05], but the difference was not significant between B_NF and BM [*p* = 0.25], and between B_NF and VPL [*p* = 0.39]. Performance accuracy in the BM condition was significantly greater than VPL and VPR [*p* < 0.05]. At T1, the performance accuracy in B_NF was significantly greater than VPL and VPR [*p* <= 0.05], but the difference was not significant between B_NF and BM [p = 0.16]. The accuracy of BM was significantly better than VPL and VPR [*p* < 0.05], while the difference between VPL and VPR was not significant [*p* = 0.32]. At T2, performance accuracy of B_NF was significantly better than VPL and VPR [*p* < 0.05]. The difference between B_NF and BM [*p* = 0.18] was not significant. Performance accuracy in the BM condition was significantly better than VPL and VPR [*p* < 0.05], but the difference between VPL and VPR was not significant [*p* = 0.69]. These results reveal that bimanual practice facilitated bimanual coordination accuracy without visual feedback during the retention test at T1 and T2.

**Figure 5.**
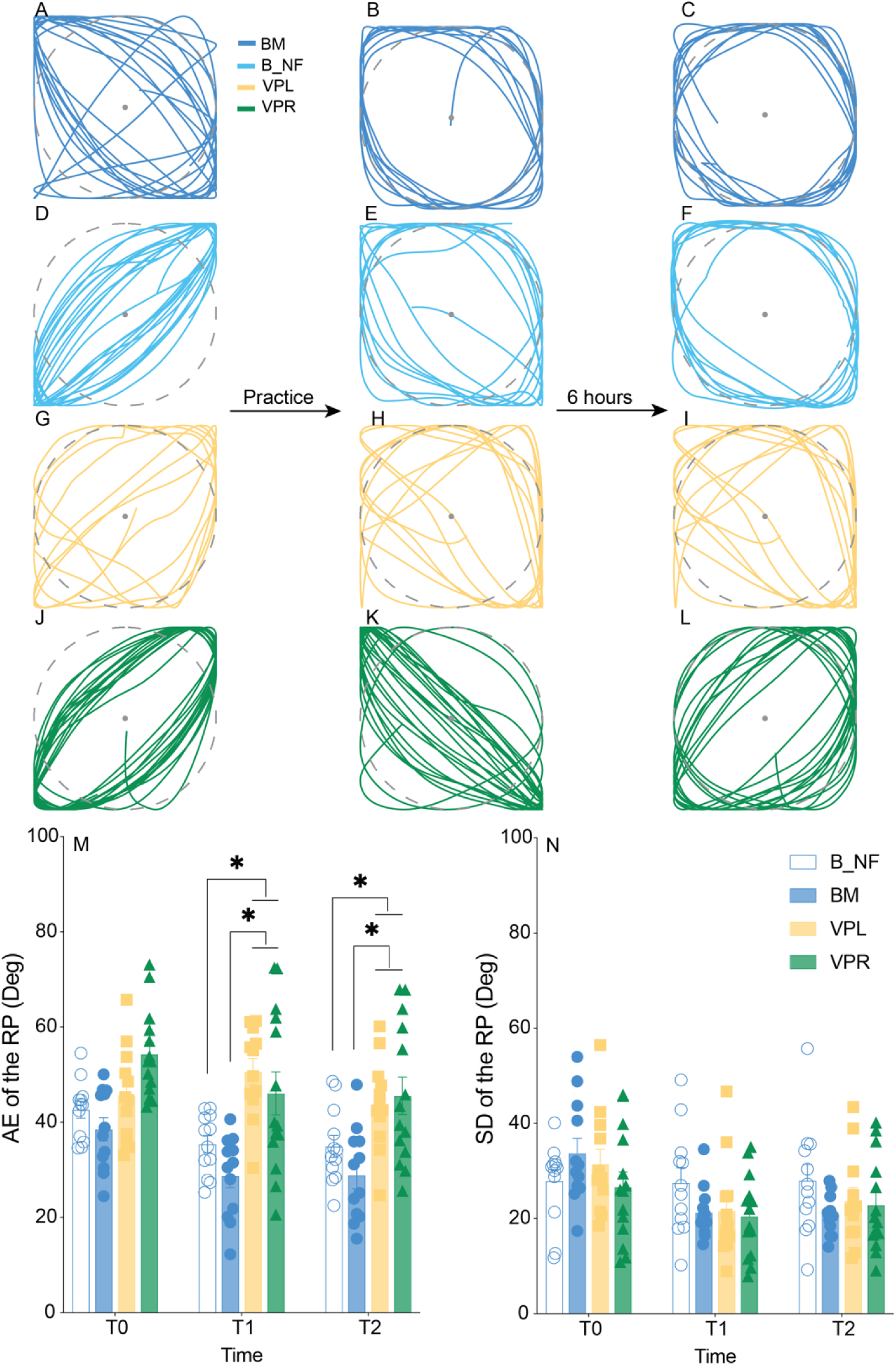
(A-L) Example performance traces for each of the conditions at each testing trial (T0, T1, T2).The AE of relative phase (M) and the standard deviation of relative phase (N) for each of the 3 testing trials (T0, T1, T2) plotted by condition plotted by condition: B_NF condition open circles (white), BM training closed circle (blue), bimanual performance after VPL training (squares yellow), and bimanual performance after VPR training (triangles green) are represented. * Indicates statistical significance (p<0.05).

The analysis of ϕSD revealed a significant main effect of Time [F (2, 92) = 20.08, p < 0.01, partial η^2^ = 0.30], and a significant Time × Condition interaction [F (6, 92) = 3.27, *p* < 0.01, partial η^2^ = 0.18] (Figure 5N). The post hoc analysis indicated an overall improvement in coordination stability, with bimanual performance being more stable at T1 and T2 compared to T0 [*p* < 0.01]. There was no significant difference in ϕSD between each condition at T0, T1, and T2. BM stability increased around the 90° coordination pattern, with variability at T1 and T2 less than at T0 [p < .01]. Coordination stability for B_NF remained unchanged over time.

## DISCUSSION

In the current investigation, we employed and modified the interlimb transfer paradigm to determine the control mechanisms and potential neural processes underlying bimanual coordination. The performance of the 90° coordination pattern improved in both BM and unimanual-VP conditions with augmented visual feedback provided during practice, indicating the representation of the given coordination pattern was developed. Following initial practice, the 90° coordination pattern acquired by one limb positively transferred to the opposite untrained limb, irrespective of the order in which limbs were practiced, and the transfer persisted after a 6-hour wakeful period. The current findings further support the notion that a given coordination pattern is represented abstractly in a task space that is independent of the effector system performing the action (26, 31–33). The discussion focuses on the nature of the shared motor skill representation and the strengths and weaknesses of plausible behavioral and neural theories of motor learning and transfer.

### Acquisition of the novel coordination pattern

Unlike intrinsic coordination patterns such as in-phase and anti-phase, the generation of the 90° coordination pattern often necessitates extensive practice lasting for several consecutive days (5, 36). The current experiment aligns with previous findings by demonstrating that providing augmented visual feedback significantly reduces the typical practice time required for mastering complex coordination patterns (1, 3, 4, 13, 37, 38, 40, 42).

Previous research suggested that visual information facilitates the quick development of the motor memory representation in extrinsic coordinates, at least in the early stages (13, 47–49). This process has been highlighted in a deafferented patient study as proprioception independent (50), where deafferented patients outperform healthy individuals when tracing with mirror-reversed visual feedback. Moreover, EEG recorded from the somatosensory cortex in healthy individuals has revealed a suppression of proprioceptive inflow in the early phase of learning during visually guided movement (48). As such, given the fact that unimanual-VP coordination is predominantly coupled through vision (43, 51, 52), the memory representation of the coordination pattern is quickly developed using the visuospatial information to support the performance of unimanual-VP coordination. Specifically, the movement of the VP and coordination errors can only be captured and analyzed by the visual system to recalibrate the kinematics of the acting limb, information that is unavailable in the proprioceptive feedback of the acting limb (32, 43).

Moreover, our results provide substantial evidence supporting the existence of bimanual and unimanual-VP memory, as the practiced skills were consolidated during the 6-hour post-training period (Figure 2A&B). Notably, the performance of each group at the 6-hour retention test remained consistent with the end-of-practice performance and was significantly improved compared to the initial performance (Figure 2A&B). Consolidation, recognized as an offline process that stabilizes initial memory, has been demonstrated in motor learning across various paradigms (53–56), including bimanual coordination (13, 40). The present experiment uniquely reveals memory savings for unimanual-VP coordination at the 6-hour retention test. Had unimanual-VP coordination undergone only short-term adaptive processes, a sharp decline in memory would be expected during the retention test after 6 hours. Consequently, we posit that consolidation has indeed occurred following unimanual-VP practice. Nonetheless, further studies using the retrograde interference paradigm would be necessary to obtain conclusive evidence for this assumption (40, 56, 57).

### Motor generalization across limbs

To date, numerous studies have explored motor generalization across various tasks (16, 21, 26, 28, 29, 31, 33, 47, 58). The success of generalizing acquired motor skills to non-practiced effectors or conditions has been demonstrated to be task-dependent and/or condition-dependent (16, 21, 47, 58). For instance, the positive generalization of the reaching direction has been observed from the dominant to the non-dominant limb in right-handed individuals during visuomotor adaptation (20). Conversely, in the context of dynamic force adaptation, this generalization occurs from the non-dominant to the dominant limb (21). Furthermore, the extent of task practice has been linked to the effect of generalization (28). Kumar and colleagues (2020) have explored generalization effects using a visuomotor adaptation paradigm (28). The adaptation process involves both fast and slow processes, with early adaptation attributed to the fast process (59, 60). Consequently, generalization is believed to stem from slower implicit processes associated with refining internal motor representations rather than the fast process involving explicit strategies (28).

However, this assumption may not apply because of the advantage of visual feedback in the current experiment, where the integrated visual feedback significantly reduces the incidental constraint on performing the given coordination pattern (41). As such, improved performance in these transfer conditions could stem from an abstract enhanced perceptual representation after brief exposure to the task rather than extended physical practice alone (61, 62). To eliminate this possibility, we conducted a control experiment where participants performed the required task conditions for two sessions separated by 6 hours without engaging in physical practice beforehand. However, results indicate that neither the augmented visual feedback alone nor short exposure to the task content provides benefit for subsequent non-practiced conditions, including during testing administered 6 hours later. Improved performance in non-practiced conditions is primarily driven by the physical practice of the initial condition (Figure 3A-D; Figure 4A&B), with the refinement of motor performance via implicit process (28). The generalizability of motor skill knowledge to non-practiced conditions remains unchanged in the 6-hour retention test. In alignment with the findings of Kumar (2020), this result underscores that the degree of generalization is directly correlated with the quality of the motor representation in the memory trace (28).

In the context of generalization between unimanual conditions, prior practice has been shown to enhance subsequent task performance, as evidenced by improved accuracy (AE of the coordination) and coordination stability (SD of the coordination) compared to the control condition (Figure 3A-D), irrespective of the hand trained first. The validity of this finding is bolstered by previous research that suggests a 90° coordination pattern acquired during inter-joint coordination within one limb can be shared to and replicated by the opposite limb (26, 31). This effector-independent characteristic linked to coordination patterns is considered an abstract representation of the spatiotemporal relationship between two coordinated components (31). The memory representation of the given coordination pattern is expected to guide the interaction between moving objects within the same environment, regardless of the physical and mechanical properties of the involved components. In the current investigation, the memory representation of the target coordination pattern acquired in each unimanual condition not only enhanced the performance of the non-practiced hand but also facilitated performance of the 90° coordination pattern in the bimanual condition (Figure 4A&B). This finding is consistent with previous research (32), but we further demonstrate that the degree of transfer to the bimanual condition is similar regardless of the hand trained first. The transfer of learning that occurs between unimanual and bimanual coordination is believed to result from the two tasks sharing common task dynamics conditioned by perceptual information and intrinsic stability (32). The change in either the perceptual information or the intrinsic stability of the task could lead to poor transfer between the two conditions (32).

It is noteworthy that bimanual coordination stability, as reflected by the SD of the coordination, was similar in both the immediate and 6-hour retest across all conditions. This seemingly counterintuitive result challenges the extent of generalization to bimanual coordination arising from unimanual-VP practice. However, a detailed analysis from a dynamic theory perspective reveals the absence of the formation of a new attractor at 90° in the control condition, as indicated by the stabilization of coordination around the intrinsic patterns of 0° and 180° (Figure 4C-F). Consequently, the observed similarity in bimanual stability is achieved but is not centered around the required 90° pattern.

Transfer tests frequently serve as a tool to elucidate the underlying mechanisms of motor control. The present transfer findings suggest potential support for the callosal access hypothesis over the dynamic dominance hypothesis concerning memory storage and sharing in governing both unimanual and bimanual coordination. The dynamic dominance hypothesis, contradicting the current results, posits distinct control mechanisms within each primary effector-control system, as evidenced by unique kinematic characteristics in movement trajectories (20, 21). Alternatively, the callosal access hypothesis proposes that memory representation governing movements of both hands is lateralized to the dominant hemisphere, shared via the corpus callosum (63), granting equal access to both hands. Previous research utilizing EEG (6), electrocorticography (ECoG) (64, 65), and fMRI (8) has provided compelling evidence suggesting lateralized memory representation governing left- and right-hand movements in unimanual and bimanual conditions. Clinical studies have revealed a greater deficit in bimanual movement planning among patients with left hemisphere damage, with those with right hemisphere damage showing no significant differences from healthy controls (24). Consequently, in the present study, it is plausible that the memory representation of the coordination pattern, acquired regardless of practice conditions, is predominantly developed, and stored in the left hemisphere. This lateralized memory representation, encoding perceptual information and task dynamics of the target coordination pattern, likely contributes to the coupling between coordinated components irrespective of the condition.

### Bimanual coordination in the absence of cursor feedback

Motor learning, as elucidated by prior research, encompasses the simultaneous development of memory representations within distinct reference systems (49). For instance, the acquisition of motor sequences initiates with the assimilation of associative responses grounded in visual stimuli. The manifestation of this learning within the visuospatial domain becomes evident through motor generalization conditions. Under these conditions, similar reactions to the same sequential stimuli can occur with the opposite hand, even though different effectors are used in the action (49, 66, 67). The establishment of memory representation, rooted in intrinsic proprioceptive feedback, enables the execution of the sequential order to be less dependent on the initial visual guidance. This execution involves corresponding effectors in the opposite hand reacting to mirror the reverse sequential order of the visual stimuli. In general, the process of motor learning entails distinct timescales in developing both spatial and motor representations, with each process being supported by different sensory information (49, 66–68).

The acquisition of a novel bimanual coordination pattern, facilitated by augmented visual feedback, often establishes a pronounced dependence on visual guidance during subsequent retesting sessions (37, 41). During post-practice, a breakdown in bimanual performance is often observed as the cursor feedback employed during the initial practice disappears (37). Subsequent research has demonstrated that the intensity of this visual guidance can be mitigated by recalibrating the relative contribution of visual and proprioceptive information throughout the learning process (1). The strength of the motor representation of the practiced bimanual coordination pattern increases as more attention is split into taking advantage of intrinsic proprioceptive feedback (1). This adjustment showcases the adaptability of the motor learning system and emphasizes the dynamic interplay between sensory modalities in the refinement of bimanual coordination patterns (69).

The present findings demonstrate the ability to produce the prescribed bimanual coordination pattern in the absence of feedback during post-practice sessions after targeted bimanual coordination training (Figure 5A). This finding appears to contradict the results reported by Kovacs and colleagues (2009), wherein they observed an immediate breakdown in the execution of bimanual 90° coordination upon the removal of the cursor (37). Their results assert a pronounced guidance effect attributed to the utilization of augmented visual feedback during bimanual coordination training, potentially compromising the utilization of proprioceptive feedback (1). Nevertheless, it is crucial to note that the duration of practice in the present experiment is twice that of Kovacs et al. (2009), potentially providing ample time and information essential for the comprehensive development of the motor representation for the specified bimanual coordination pattern. In the current investigation, the quality of bimanual performance without the cursor feedback tends to be no different from the bimanual performance facilitated by the visual feedback. During the 6-hour retention test, this performance persists (Figure 5A), underscoring the role of consolidation in stabilizing the initial motor representation associated with the given bimanual practice (13, 40).

The absence of cursor feedback has led to impaired bimanual performance after practice in both unimanual conditions (Figure 5A, I-N), underscoring the pivotal role of visual information in facilitating unimanual-VP synchronization (32, 43, 51, 52). This finding may stem from the lack of intrinsic proprioceptive feedback accounting for the movement kinematics of the opposite hand during unimanual practice, hindering the development of bimanual memory in the intrinsic coordinate (1). The given proprioceptive information in unimanual practice predominantly supports the recalibration of muscle dynamics in the acting limb to orchestrate the magnitude and speed of the moving VP.

As such, the findings of the current investigation underscore the need for extensive bimanual practice after acquiring the coordination pattern to strengthen the bimanual memory through implicit calibration using intrinsic proprioceptive feedback. In addition, these results provide an example in agreement with the notion that generalization may be affected by the way in which the memory is encoded and consolidated (47, 70). In this context, our findings illustrate that memory representation formed exclusively in the extrinsic coordinate fails to contribute to subsequent performance in the intrinsic coordinate when salient visual information is absent. Consistent with previous literature, these results highlight the simultaneous occurrence of error-based movement correction rooted in information derived from both extrinsic and intrinsic references during bimanual coordination (34, 71, 72).

### Neural substrates encoding effector-independent representations

It has been long suggested that the primary motor cortex (M1) is involved in motor learning, mediating online and offline processes (73). Previous studies have consistently demonstrated that disruption to M1, occurring before (74), during (75), and after (76) motor skill practice, results in impaired motor performance at the retention test. In bimanual coordination tasks, increased M1 excitability in left M1 (of right-handed individuals) emerges during (12) and after (13) practice. Notably, the degree of left M1 excitability exhibits a positive correlation with the task’s difficulty, with more complex 90° bimanual coordination inducing greater excitability than in-phase and anti-phase bimanual coordination patterns (12). These findings support the involvement of left M1 in the performance of bimanual coordination, implying the possibility that bimanual coordination is lateralized to the dominant hemisphere. The left hemisphere houses task-relevant information utilized in controlling ipsilateral effector movement via interhemispheric connections (77, 78). This interhemispheric mechanism plays a crucial role in inter-limb transfer and bimanual coordination, potentially mediated by M1 activity. For example, a prior investigation illustrated that the modulation of left M1 excitability through transcranial direct current stimulation (tDCS) enhances skill transfer rates to the untrained hand post-adaptation in a dynamic force field (79). This augmented generalization might result from M1 plasticity, influencing the motor learning pattern (79).

Interhemispheric inhibition (IHI) regulates communication between M1 regions and is mediated by the corpus callosum (CC) (77, 80). Previous studies suggest that corticospinal output to the resting limb is influenced by the active limb, mediated by IHI (77, 81), hinting at a potential influence of IHI modulation on generalization to the non-practiced limb. A recent study found a positive correlation between individual learning-related changes in IHI and the degree of generalization (58). Reduced IHI, indicating enhanced access to contralateral hemisphere information, is associated with improved performance in the non-practiced limb (58). These results highlight the role of the CC in facilitating interhemispheric information sharing and indirectly support M1’s involvement in generalization. Given the CC’s established role in synchronizing bimanual actions (6, 78, 81), it is reasonable to expect its crucial involvement in generalizing the acquired coordination pattern studied here.

The left PPC may contribute to limb-independent generalization, as suggested by previous research on abstract motor plans and memory structures (82–84). Elevated cortical activity in the left PPC during motor sequence learning is associated with associative responses to visuospatial cues (84). Furthermore, cathodal tDCS applied to the left PPC, but not the right, impedes limb-independent generalization (28), indicating potential lateralization in storing limb-independent representations. fMRI data indicate that a left parietal-to-(pre)motor network predominantly activates during 90° bimanual coordination, suggesting encoding by cortical regions in the left hemisphere, such as M1 and PPC (8). However, our study does not investigate specific neural regions responsible for unimanual and bimanual coordination using neurophysiological and non-invasive approaches like TMS, EEG, and fMRI. Thus, direct evidence of lateralization in cortical activations is lacking. Behavioral results imply potential lateralization in bimanual coordination, warranting further investigation into cortical activation and interhemispheric communication involved in both types of coordination in our next phase of the investigation.

This study stands out as the first to demonstrate that unimanual practice, irrespective of the hand trained first, can indeed benefit the performance of bimanual coordination subsequently by structuring a task that minimizes discrepancies in movement patterns between unimanual and bimanual actions. This result holds significant implications for the design of future rehabilitation approaches aimed at restoring bimanual functions. Many daily tasks rely on the ability to coordinated both arms together (85, 86). The inability to coordinate bilateral actions significantly impacts the quality of life among individuals recovering from unilateral stroke (87). The findings from this study suggest a potential avenue for restoring motor functions in the paretic limb and improving bimanual coordination by training the unaffected limb in individuals with unilateral stroke.

## SUMMARY

In this article, we aimed to discern the individual contributions of each limb in the acquisition of a novel bimanual coordination pattern. The results discussed above indicate that each limb plays a role in the evolution of the representation associated with the 90° coordination pattern. The pattern acquired through each limb can be successfully reproduced in bimanual coordination when the same visual feedback is provided. However, the inability to replicate 90° bimanual coordination after practicing each limb without the aid of the cursor underscores the pivotal role of the interactive coordination between acting limbs in shaping the intrinsic motor representation of bimanual coordination. The noteworthy findings from this experiment offer exciting possibilities for designing specific therapeutic approaches aimed at restoring bimanual function in stroke survivors, leveraging the utilization of available information within their sensorimotor system.

## GRANTS

This work has received no support from any grants.

## ACKNOWLEDGMENTS

The authors would like to thank the participants for volunteering their time.

## AUTHOR CONTRIBUTIONS

Y.W., Y.L., D.L.W., J.J.B., and D.M.K. designed research; Y.W., and M.M.W. performed research; Y.W., J.J.B., and D.M.K. analyzed data; and Y.W., M.M.W., Y.L., D.L.W., M.S., J.J.B., and D.M.K. wrote the paper.

## Data availability statement

The data that support the findings of this study are available from the corresponding author upon reasonable request.

